# *Mycobacterium tuberculosis* Δ*lsr2* Vaccine Enhances Protection Against Tuberculosis

**DOI:** 10.64898/2026.07.07.737086

**Authors:** Jacqueline Watt, Jun Liu

## Abstract

Tuberculosis (TB) has been a leading cause of death from a single infectious agent for decades. Bacille Calmette-Guérin (BCG) remains both the primary TB vaccine strategy and the oldest vaccine in circulation, with severe limitations in adult populations. Recently, live attenuated vaccine strategies, or generating safe strains of *Mycobacterium tuberculosis* (*Mtb*) through genetic engineering, have shown considerable promise. We previously developed an attenuated strain of *Mtb* lacking the nucleoid-associated protein Lsr2 (Δ*lsr2*) which is also phthiocerol dimycocerosates (PDIM) deficient and induces an immune response that represents an intermediate stage between the parental *Mtb* strain and BCG. In this study we examined the immune response of Δ*lsr2* vaccinated mice in comparison to BCG and found a substantially stronger CD4 and CD8 T cell responses from Δ*lsr2* vaccinated mice. Complementary, we conducted *Mtb* protection studies in Δ*lsr2* and BCG vaccinated mice and guinea pigs, where we found that Δ*lsr2* provided superior protection in both animals. This improved protection is shown with reduced bacterial burden and improved organ pathology in the lungs and spleen. Taken together, our work shows Δ*lsr2* serves as a promising vaccine candidate for continued preclinical development.

## Introduction

Tuberculosis (TB), caused by the bacterium *Mycobacterium tuberculosis* (*Mtb*), remains one of the world’s most pressing infectious diseases, with approximately ten million new cases and over one million deaths annually ^1^. Despite over a century of widespread Bacille Calmette-Guérin (BCG) vaccination, global TB incidence has remained largely unchanged. While BCG effectively protects against severe pediatric TB, its efficacy against pulmonary TB in adolescents and adults is highly variable ^2^. The scale of the TB epidemic, coupled with an estimated 1.7 billion individuals harboring latent infection, underscores the urgent need for more effective vaccines ^1^.

One approach to improving TB protection in adults is the development of subunit booster vaccines, which deliver selected antigens to enhance protective immunity initially conferred by BCG. A notable example is the Modified Vaccinia Ankara virus expressing antigen 85A (MVA85A) vaccine, designed to induce CD4 and CD8 T cell responses against antigen 85A (Ag85A), a major secreted protein of *Mtb* involved in cell wall biosynthesis and strongly recognized by the host immune system ^3^. Ag85A is present in all BCG strains, thus the MVA85A strategy aimed to boost and prolong BCG-induced protection ^4^. Early animal vaccine studies were promising, showing induction of appropriate T cell responses and modest improvements in protection, however, MVA85A ultimately failed to enhance protection in human trials ^3,4^.

Another approach involves the addition of adjuvants to subunit vaccines to illicit better immune responses, such as the recent M72/AS01ₑ vaccine candidate. M72/AS01ₑ delivers two *Mtb* proteins, Mtb32A and Mtb39A, implicated in pathogen survival and virulence along with the GSK adjuvant AS01ₑ ^5^. Initial clinical trials indicate M72/AS01ₑ confers protection of approximately 50% in *Mtb* infected adults, however it is important to note this was designed for Prevention of Disease (POD) but not for Prevention of Infection (POI) ^5^.

Live attenuated vaccines represent a complementary strategy, aiming to more closely mimic natural infection and elicit broad, durable immune responses. The *Mtb* complex (MTBC) has been associated with humans for over 50,000 years, consequently MTBC has become highly human-adapted and is only able to efficiently transmit, establish and maintain infection cycles in humans ^6^. BCG is a highly attenuated strain derived from *Mycobacterium bovis*. Contrary to previous belief, humans likely did not originally acquire tuberculosis from *M. bovis* during early cow domestication; phylogenetic analysis has revealed certain *Mtb* lineages are more ancestral than *M. bovis* ^6^. BCG has 25 identified ‘Regions of Difference’ (RDs), many of which encode virulence-associated antigens that are common targets for new TB vaccine candidates, such as RD1 which encodes culture filtrate protein 10 (CFP-10) and early secretory antigenic 6-kDa (ESAT-6) ^7,8^. Moreover, BCG has lost 23% of the experimentally confirmed MTBC T cell epitopes that are essential to the host immune response ^9^. Our immunological profiling work in mice has demonstrated distinct immune signatures between BCG and *Mtb* across diverse lung immune cell populations ^10^. These observations suggest that attenuating *Mtb* may generate protective immune responses unattainable with BCG-based vaccine strategies alone ^10^.

One promising live-attenuated vaccine candidate is MTBVAC, which harbors deletions in the virulence genes *phoP* and *fadD26* ^11^. MTBVAC has shown superior immunogenicity and protective efficacy compared with BCG in animal models, eliciting strong immunogenicity and an acceptable safety profile ^11^. Phase 1 clinical trials have confirmed MTBVAC’s safety and immunogenicity in adults ^12^. Further Phase 2 and Phase 3 clinical trial evaluations of MTBVAC are currently ongoing.

We have previously characterized a strain of *Mtb* with the loss of Lsr2 (Δ*lsr2*), a nucleoid-associated protein that functions as a global transcriptional repressor ^13–16^. We have also shown its severe attenuation in mice and reported that this strain is phthiocerol dimycocerosates (PDIM) deficient, most likely due to spontaneous mutation from *in vitro* passage ^10^. Interestingly, the second gene deletion in MTBVAC, *fadD26*, is also responsible for PDIM biosynthesis ^17^. We also previously described our systematic profiling of lung immune cells in mice following inoculation with *Mtb*, Δ*lsr2*, or BCG using cytometry by Time-Of-Flight (CyTOF) where we observed the immune response elicited by Δ*lsr2* represented an intermediate stage between the parental *Mtb* strain and BCG, with respect to bacterial burden, immune cell infiltration, T cell activation and exhaustion markers ^10^.

Building on the precedent set by MTBVAC, our Δ*lsr2* strain leverages rational genetic attenuation to balance safety and immunogenicity. Here we evaluated Δ*lsr2*’s induced T helper cell type I (Th1) cytokine response to *Mtb* including IFNγ and TNF, which are well documented to be critical for control of *Mtb* in both humans, mice and non-human primates ^18^. We also investigated Δ*lsr2*’s protective efficacy against *Mtb* infection in female C57BL/6 mice and guinea pigs. Δ*lsr2* induced a stronger immune response of Th1 cytokines and improved protection relative to BCG, demonstrating the feasibility of genetically attenuated *Mtb* vaccines as a next-generation strategy to achieve durable protection against TB.

## Results

### Δlsr2 induces higher levels of antigen specific CD4 and CD8 T cells than BCG

To investigate whether Δ*lsr2* induces improved immunogenicity over BCG, we aerosol inoculated a group of six female C57BL/6 mice with either ∼100 colony forming units (CFU) of Δ*lsr2* or BCG using the Glass-Col Inhalation Exposure System (Figure 1a). After 6 weeks, mice were humanely euthanized, and the lungs and spleen processed for intracellular cytokine staining (ICS) and flow cytometry analysis. Δ*lsr2* induced a much higher percentage of antigen specific CD4 T cells expressing IFNγ and/or TNF (∼50-100-fold; p < 0.05) and CD8 T cells expressing IFNγ (∼10-fold; p < 0.05) in the lungs than BCG (Figure 1b). In the spleen Δ*lsr2* induced a ∼10-fold higher percentage of IFNγ+ CD4 T cells (p < 0.05; Figure 1c).

**Figure 1.**
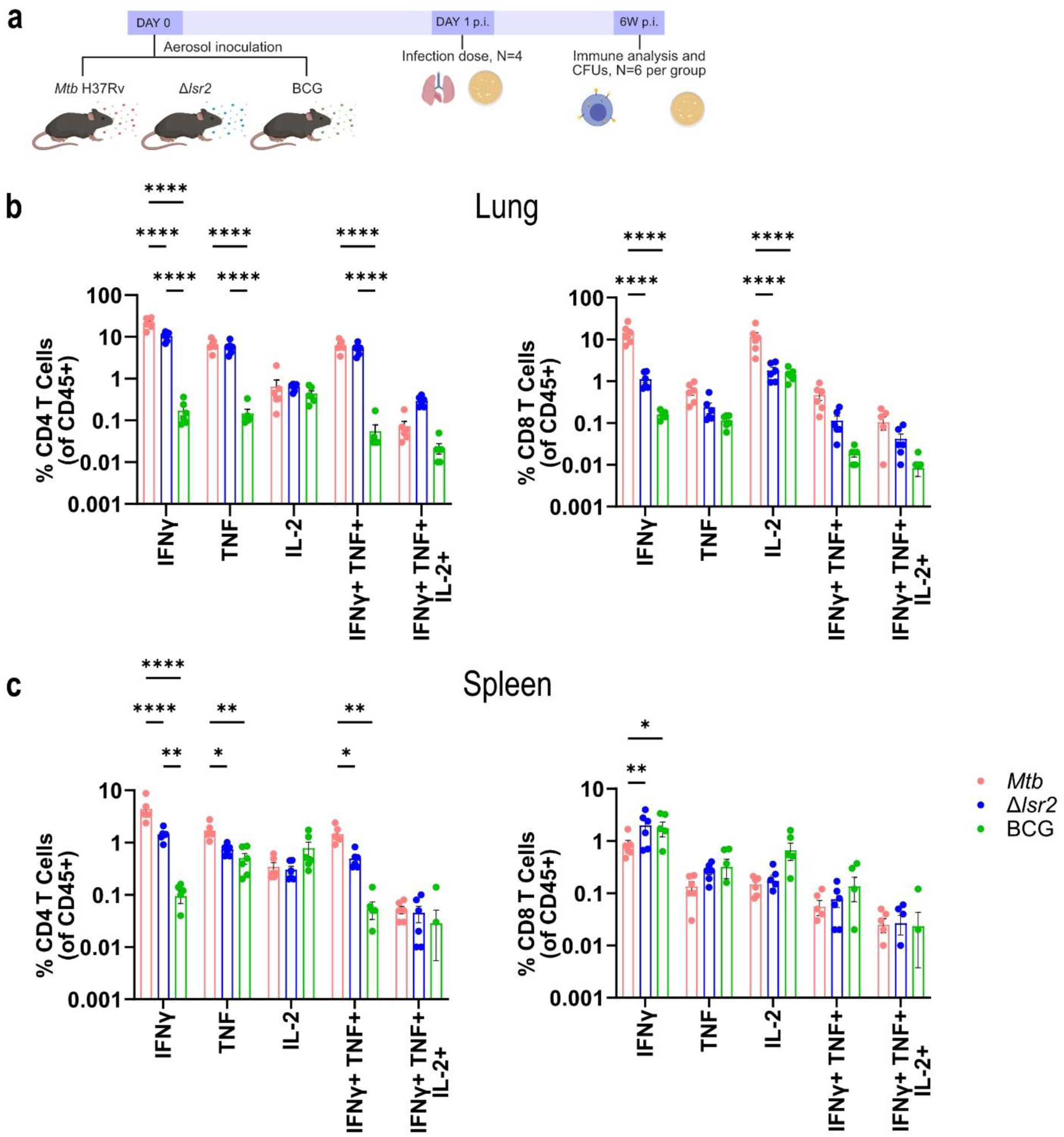
Δ*lsr2* induces a strong IFNγ and TNF response in CD4 and CD8 T cells similar to the parental strain *Mtb* H37Rv. a) Schematic representation of experimental design. 6 C57BL/6 mice per group were aerosol inoculated with *Mtb* H37Rv, Δ*lsr2* or BCG-Japan (∼100 CFU/lung). At 6 weeks p.i. lung and spleen samples were processed into single cell suspensions and treated for 24 hours with *Mtb* lysate to determine differences in the production of Th1 cytokines IFNγ, TNF and IL-2. The percentage of CD4 and CD8 T cells producing Th1 cytokines in the b) lungs and c) spleen was determined. Data presented as mean ±SEM. Statistical significance was determined by two-way ANOVA with Tukey’s multiple comparisons test (*, p<0.05; **, p<0.01; ***, p<0.001; ****, p<0.0001; ns = not significant).

We next sought to compare the induced immune response of Δ*lsr2* to that of an *Mtb* infection by using a third group of C57BL/6 mice that were aerosol immunized with the parental strain *Mtb* H37Rv (Figure 1a). We previously found Δ*lsr2* to be highly attenuated with ∼3 log_10_ lower bacterial burden compared to *Mtb* H37Rv ^10^, however, here we observed that Δ*lsr2* induced only a 2-fold lower percentage of IFNγ and/or TNF expressing CD4 T cells in the lungs compared to *Mtb* (Figure 1b). A similar result was obtained in the spleen (Figure 1c). Taken together, these results suggest that deletion of *lsr2* in *Mtb* reduces virulence while preserving immunogenic potential, making it an ideal candidate for live-attenuated vaccine development.

### Δlsr2 confers better protection over BCG in C57BL/6 mice

To evaluate the protective efficacy of Δ*lsr2* compared to BCG we first conducted a mouse model infection experiment. We intranasally vaccinated groups of six C57BL/6 mice with 5×10^5^ CFU of Δ*lsr2* or BCG or PBS as a control for 8 weeks and then aerosol challenged with *Mtb* H37Rv using the Glass-Col Inhalation Exposure System (Figure 2a). The bacterial burden in the lungs was determined at 3-, 5-, and 8-weeks post infection (p.i.) and in the spleen at 5- and 8-weeks p.i. We found that intranasal vaccination of mice with Δ*lsr2* elicits protection better than BCG to *Mtb* infection in the lungs with ∼0.5 log_10_ lower CFU at week 5 (p < 0.05) and ∼0.8 log_10_ lower CFU at week 8 (p < 0.0001; Figure 2b). In the spleen, Δ*lsr2* vaccinated mice had ∼0.4 log_10_ lower bacterial burden at 5 weeks p.i. (p < 0.05) and ∼0.2 log_10_ lower bacterial burden at 8 weeks p.i. (p = n.s.; Figure 2c).

**Figure 2.**
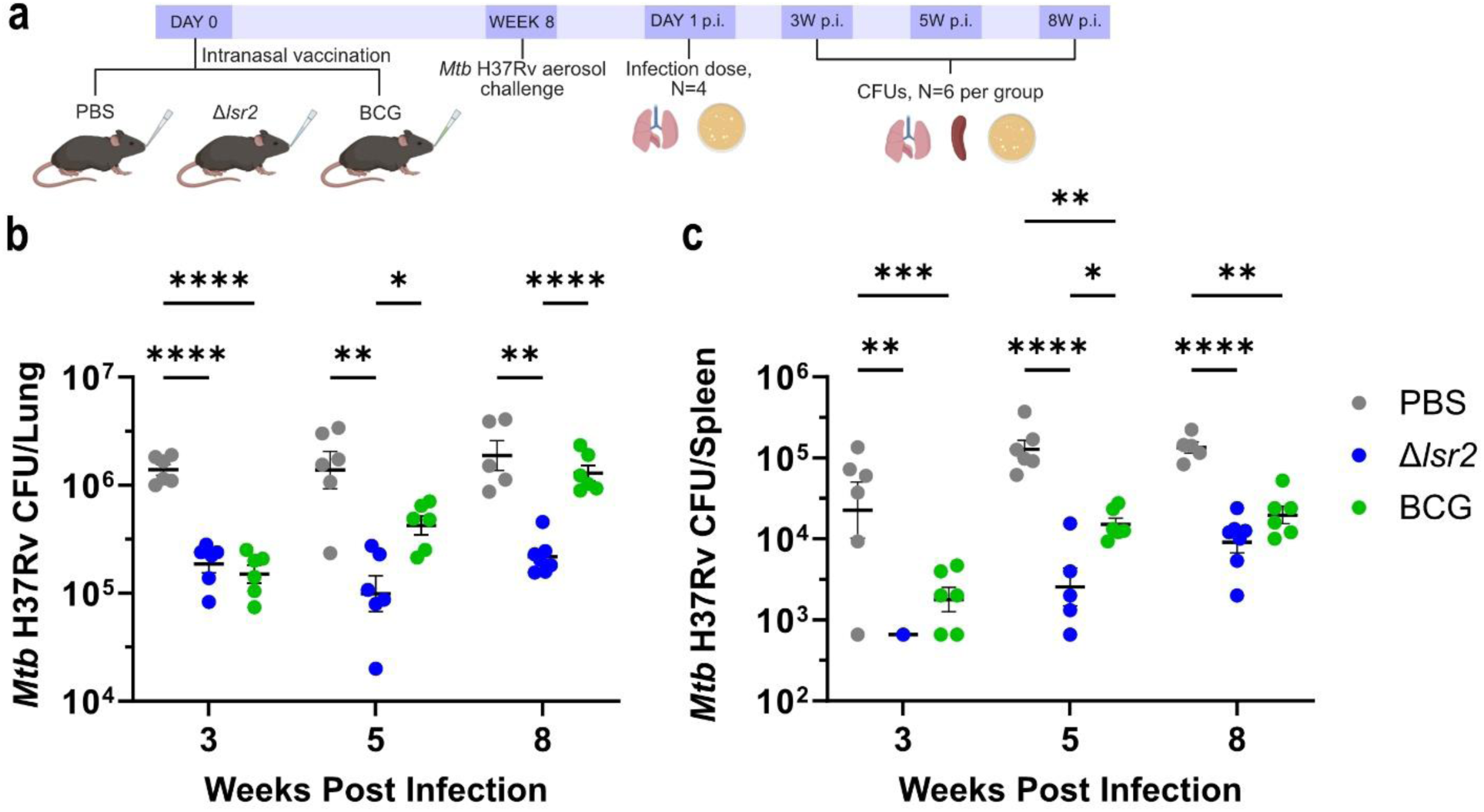
Δ*lsr2* improves protection to *Mtb* infection in C57BL/6 mice. a) Schematic representation of experimental design. 6 C57BL/6 mice per group were intranasally vaccinated with 5×10^5^ CFU of Δ*lsr2* or BCG-Japan in 20μL PBS. After 8 weeks, all mice were aerosol challenged with *Mtb* H37Rv (∼100 CFU/lung). *Mtb* H37Rv CFUs in the b) lungs and spleen c) was determined at 3-, 5-, and 8-weeks p.i. Data presented as mean ±SEM. Statistical significance was determined using a REML mixed effects model and Tukey’s multiple comparisons test (*, p<0.05; **, p<0.01; ***, p<0.001; ****, p<0.0001; ns = not significant).

### Δlsr2 displays high dissemination to the lymph nodes in C57BL/6 mice

A major bottleneck for live TB vaccines’ efficacy is delayed antigen presentation and a following delayed induction of T cell immunity ^19^. Optimal priming of T cell responses requires transport of *Mtb* infected dendritic cells from the lungs to the lymph nodes where *Mtb* antigens are then presented to naïve CD4 T cells ^19^. However, this process is very inefficient during an *Mtb* infection. One reason for this is that infected dendritic cells do not efficiently present antigens directly to naïve T cells; instead, once in the lymph node, infected dendritic cells must first pass *Mtb* antigens to uninfected bystander dendritic cells that then present antigens to naïve CD4 T cells ^20,21^. Griffiths et al. showed this deficiency can be overcome by delivering exogenously primed activated dendritic cells into lungs of vaccinated mice which led to rapid CD4 T cell activation as well as increased alveolar macrophage activation ^19^. VPM1002 used a different approach to target this same limitation, with a urease-deficient recombinant BCG that expresses listeriolysin, a hemolysin produced by *Listeria monocytogenes*, which causes leakage of BCG into the cytosol, thereby increasing early antigen presentation ^22^.

While the exact mechanism of how Δ*lsr2* provides protection remains unknown, we observed that the ratio of Δ*lsr2* CFU in the lung to the lymph node after 8 weeks of infection (Figure 3a) is ∼63% (LN CFU/lung CFU), peaking at ∼100% at 4 weeks p.i. (Figure 3b-c). This is greatly increased from a ratio of ∼3% with *Mtb* after 8 weeks of infection (p = 0.029), with the ratio for *Mtb* being the lowest at 4 weeks p.i. at ∼0.04% (Figure 3b-c). Our previous work described a highly dynamic and heterogenous dendritic cell response in Δ*lsr2* inoculated mice compared with BCG inoculated mice ^10^. Taken together, these observations could indicate antigen presentation is increased during an Δ*lsr2* infection and that more Δ*lsr2* bacilli are being transported from the lung into the lymph nodes early on during infection, potentially resulting in the strong induced immune response despite the significant attenuation of Δ*lsr2*.

**Figure 3.**
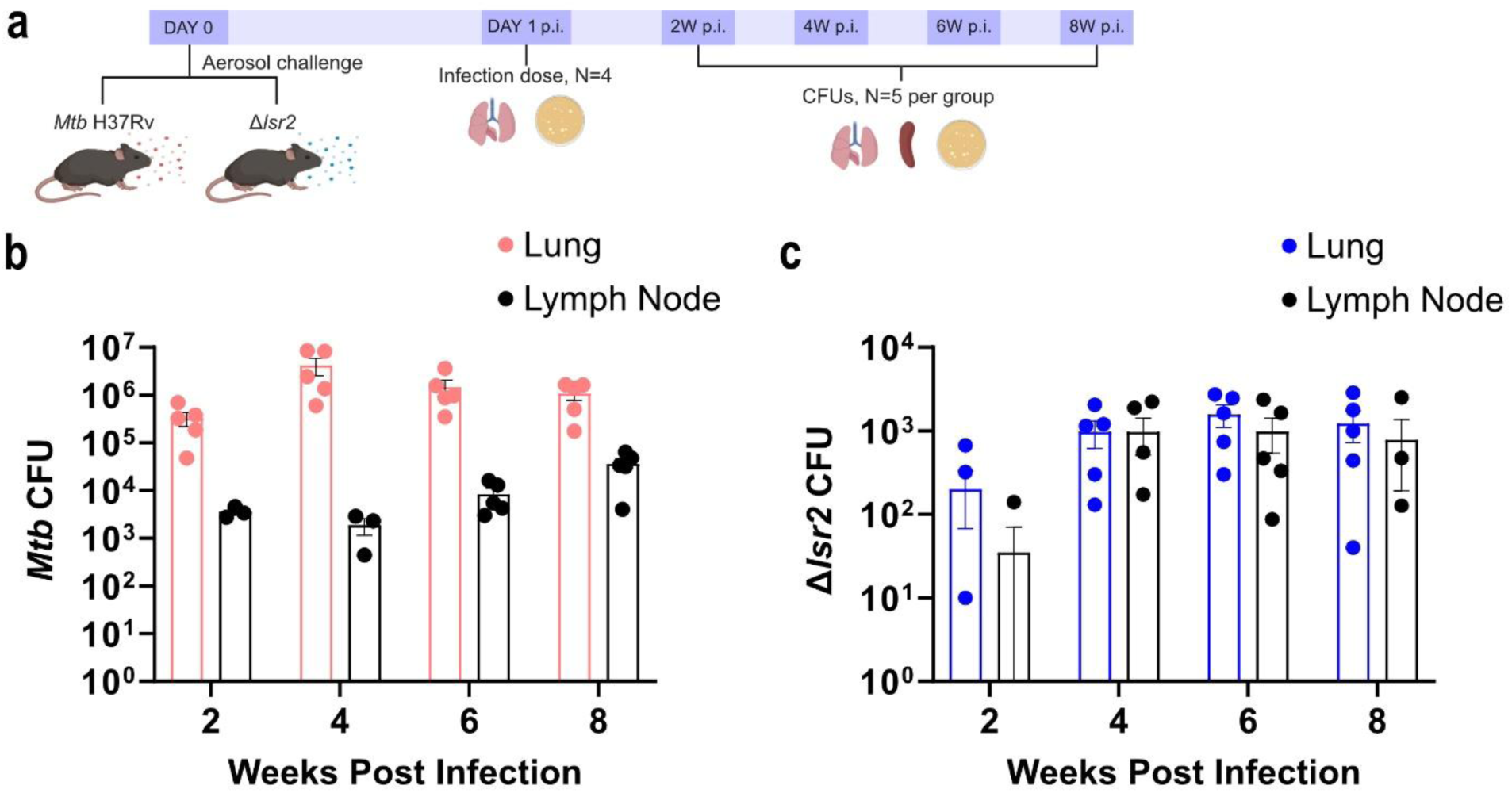
Δ*lsr2* has a high percentage of bacilli transport to the lymph nodes. a) Schematic representation of experimental design. 5 C57BL/6 mice per group were aerosol challenged with *Mtb* H37Rv or Δ*lsr2* (∼100 CFU/lung). b) *Mtb* CFU in the lungs and lung draining lymph node was determined at 2-, 4-, 6- and 8-weeks p.i. c) Δ*lsr2* CFU in the lungs and lung draining lymph node was determined at 2-, 4-, 6- and 8-weeks p.i. Data presented as mean ±SEM.

### Δlsr2 vaccination improves protection and reduces TB pathology in guinea pigs

Guinea pigs are considered a stringent test of *Mtb* vaccine effectiveness as they are highly susceptible to *Mtb* infection and develop similar symptoms to humans, such as weight loss and decreased pulmonary function ^23–26^. To further validate the efficacy of vaccination with Δ*lsr2*, we intranasally vaccinated groups of six female guinea pigs with 5×10^5^ CFU of Δ*lsr2*, BCG or PBS as a control for 8 weeks (Figure 4a). At 8 weeks post vaccination, the guinea pigs were aerosol challenged with ∼5000 CFU/lung of *Mtb* H37Rv using the Glass-Col Inhalation Exposure System. Bacterial burden in the lungs and spleen was monitored at various time points up to 7 weeks p.i. Guinea pigs were humanely euthanized once they reached the humane or experimental endpoint. 5 guinea pigs reached humane endpoint, including four in the PBS group at days 40, 42 and two on day 43 p.i., and one in the Δ*lsr2* group at day 40 p.i. Δ*lsr2* vaccinated guinea pigs had trending lower CFUs in the lungs after 7 weeks p.i. with ∼0.5 log_10_ lower CFU than BCG (Figure 4b). In the spleen, Δ*lsr2* vaccinated animals had significantly reduced bacterial burden compared to sham controls (p = 0.014) and ∼1.2 log_10_ lower CFU compared to BCG at 7 weeks p.i. (Figure 4c)

**Figure 4.**
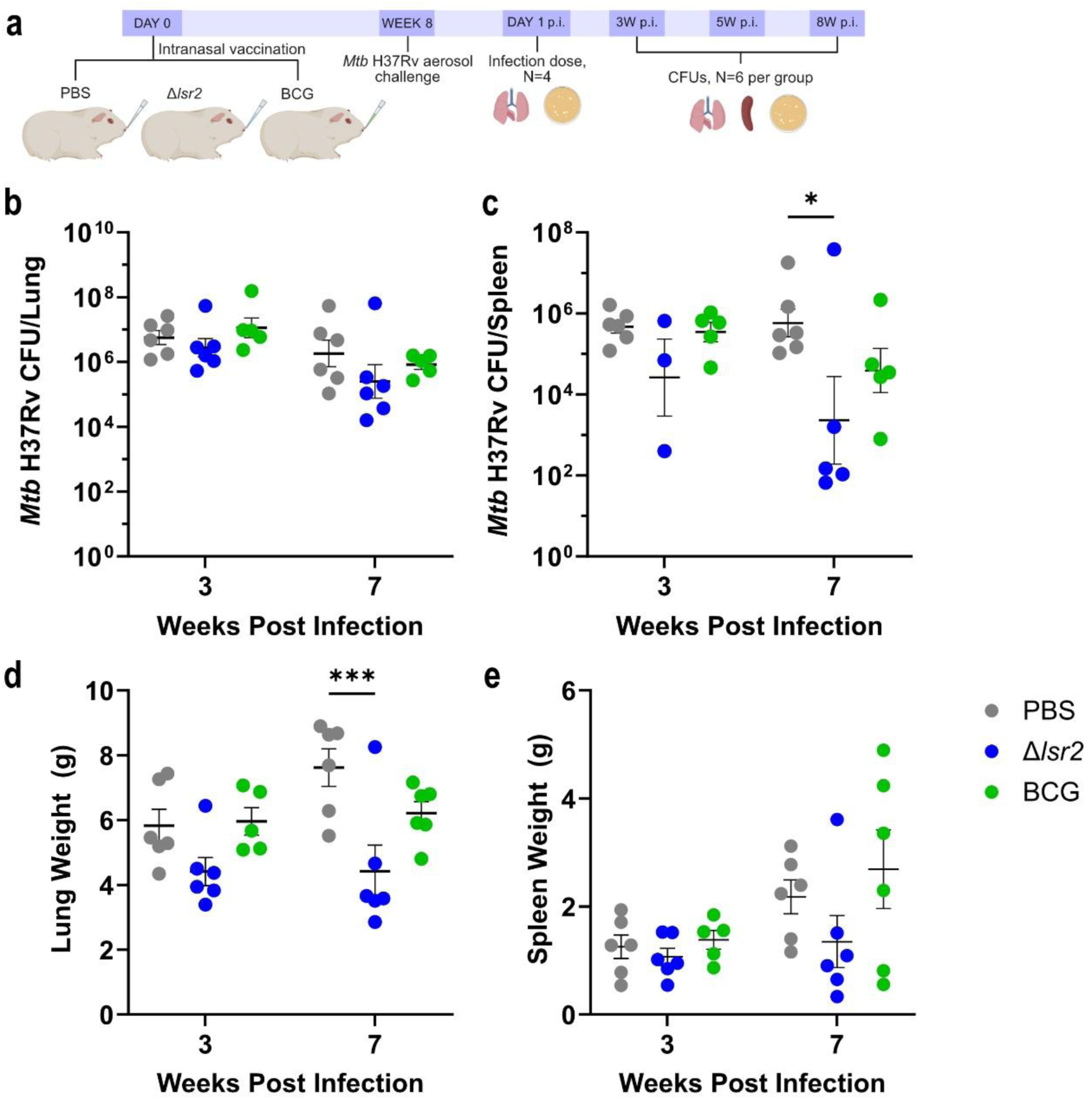
Δ*lsr2* reduces *Mtb* burden in guinea pigs. a) Schematic representation of experimental design. 6 Hartley guinea pigs per group were intranasally vaccinated with 5×10^5^ CFU of Δ*lsr2* or BCG-Japan in 20μL PBS. After 8 weeks, all animals were aerosol challenged with *Mtb* H37Rv (∼5000 CFU/lung). *Mtb* H37Rv CFUs in the b) lungs and c) was determined at 3- and 7-weeks p.i. d) Lung and e) spleen organ weights were measured at 3- and 7-weeks p.i. Data presented as mean ±SEM. Statistical significance was determined by two-way ANOVA with Tukey’s multiple comparisons test (*, p<0.05; **, p<0.01; ***, p<0.001; ****, p<0.0001; ns = not significant).

Of note, there was one non-responder outlier in the Δ*lsr2* vaccinated group with very high CFU >7.8 log_10_ in the lung, which is the previously mentioned guinea pig that reached the humane endpoint before the experimental endpoint. After removing the outlier and repeating the statistical analysis, Δ*lsr2* vaccinated animals had significantly reduced bacterial burden in the lungs (p = 0.007) compared to sham controls and ∼ 1 log_10_ lower CFU compared to BCG after 7 weeks p.i. (Supplementary Figure S1).

We further analyzed the guinea pig lungs and spleen using histological analysis and organ weight. Increased organ weights of lungs and spleen in *Mtb* infected guinea pigs is frequently reported and has been associated with more severe disease ^27,28^; conversely reduced organ weights in vaccinated guinea pigs have been associated with increased vaccine-mediated protection to *Mtb* infection compared to sham controls ^29^. Consistent with our histology results, we found that guinea pigs vaccinated with Δ*lsr2* had significantly reduced lung weights (p < 0.001; Figure 4d) and lower spleen weights (p = n.s.) after 7 weeks (Figure 4e).

Lung histological analysis of at least two randomly selected guinea pigs per vaccination group at 3- and 7- weeks p.i. revealed the extent of tissue damage in the caudal lobe was greatest in the unvaccinated animals and least extensive in Δ*lsr2* vaccinated animals at both time points. Unvaccinated animals (Figure 5a and d) and BCG vaccinated animals (Figure 5c and f) had partial to extensive consolidation, whereas Δ*lsr2* vaccinated animals appeared to have healthier tissue in the caudal lobe (Figure 5b and e) at both 3 weeks p.i. and 7 weeks p.i. Taken together our results suggest that Δ*lsr2* provides superior protection to *Mtb* infection over BCG in the stringent guinea pig infection model.

**Figure 5.**
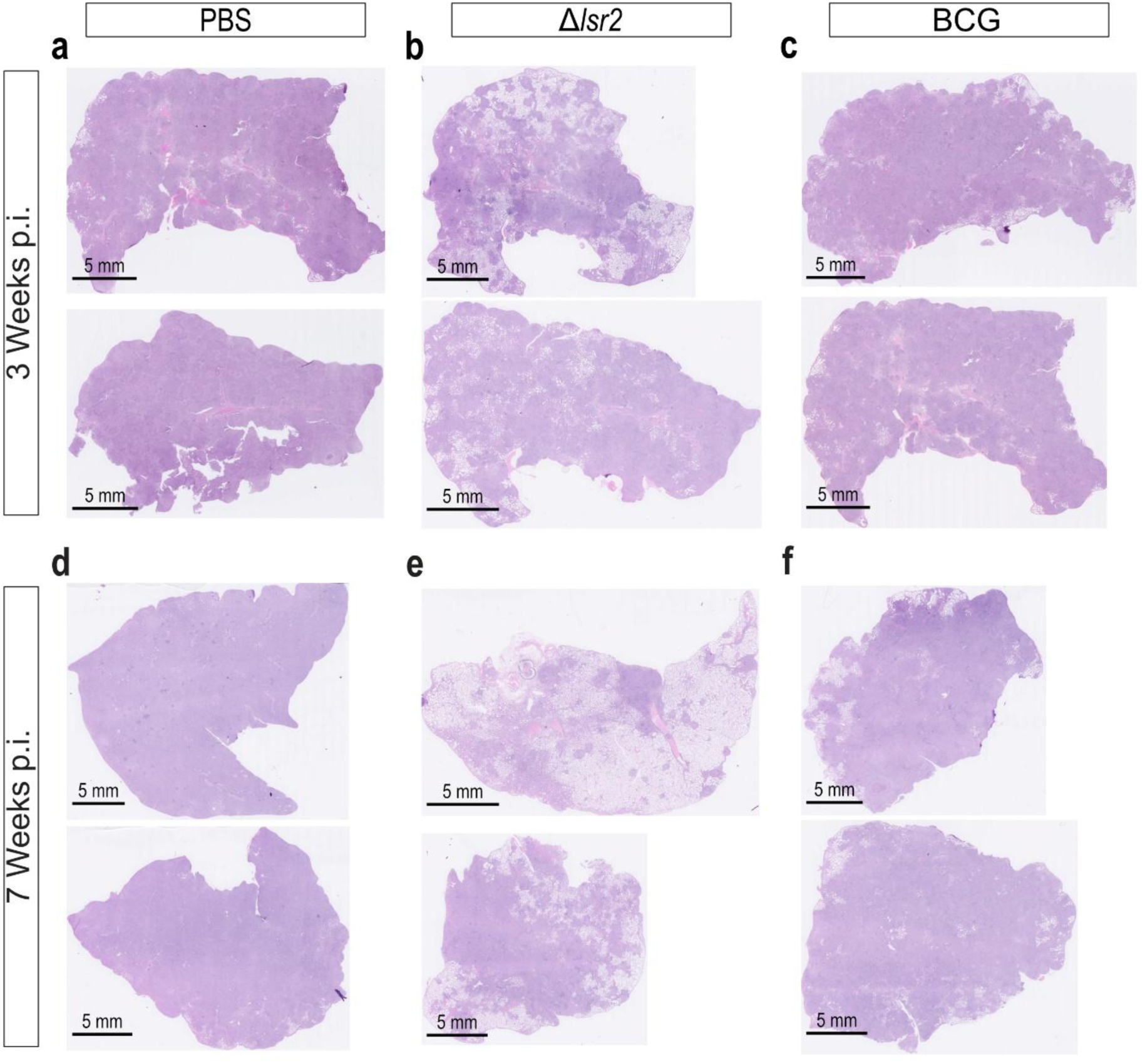
Δ*lsr2* reduces the lung pathology of guinea pigs infected with *Mtb*. Two guinea pigs from each vaccination group were randomly chosen for histological analysis. These included two guinea pigs from the a, d) PBS sham control group; b, e) two guinea pigs from the BCG group; and c, f) two guinea pigs from the Δ*lsr2* group. Histological analysis was performed at a-c) 3 weeks p.i. and at d-f) the experimental end point of 7 weeks p.i. For each guinea pig a section of the left caudal lobe was analyzed by H&E staining.

## Discussion

A large variety of strategies to improve TB vaccines are currently in development and range from subunit to live-attenuated vaccines for the purpose of preventing TB infection, TB disease or improving the outcome of TB disease treatment. As of 2025 there are 18 vaccines in clinical development, increasing from 15 in 2024, with six in Phase 3 trials ^30^. While MTBVAC remains the only live attenuated *Mtb* vaccine candidate in clinical development for prevention of TB disease, its success to date supports the idea that rational attenuation of *Mtb* specific genes could be a viable strategy to design new candidates for live vaccines against TB ^31^.

Here, my data describes Δ*lsr2* as a novel live attenuated vaccine candidate for TB. Lsr2 regulates multiple virulence-associated genes, modulates lipid metabolism, and is required for persistent infection, making it an attractive target for rationale genetic attenuation ^15,32^. Δ*lsr2* effectively provided protection from *Mtb* infection in both mice and guinea pigs better than BCG and drastically increased the level of IFNγ induced in both CD4 and CD8 T cells compared to BCG.

We observed the level of Δ*lsr2* CD4 T cell induced IFNγ was ∼100-fold greater than BCG and was more similar to the amount of induced IFNγ by *Mtb* (∼ 2-fold less). The level of BCG-induced IFNγ in this study is comparable to our previous work ^29^. T cell derived IFNγ has long been considered important in the adaptive immune response to *Mtb* infection. Genetic studies have shown that humans deficient in IFNγ are highly susceptible to TB ^33^. IFNγ induces anti-bacterial activity and restricts *Mtb* replication in the lung ^34–36^, and there is also evidence that CD4 T cell derived IFNγ correlates to protection in mouse models ^29,37^. However, a lot of emerging work is highlighting the relative importance of CD4 T cells to protection and that they are only one cell-type a part of the critical response to *Mtb* infection ^18^.

We also found that Δ*lsr2* induced a significantly higher amount of IFNγ producing CD8 T cells compared to BCG (∼10-fold). Along with CD4 T cells, CD8 T cells have also been shown to be important in protection against *Mtb* ^22,23,38,39^. While the precise role of CD8 T cells in TB immunity remains unclear, it likely involves their ability to synthesize IFNγ as well as their cytolytic mechanisms and ability to directly kill *Mtb* through the production of granulysin ^18,40,41^. Taken together this data potentially indicates an association between protection by Δ*lsr2* and T cell derived IFNγ. More in depth immune profiling of vaccine induced immunity at various timepoints pre and post *Mtb* challenge will provide greater insight into potential correlates of protection.

Guinea pigs are highly susceptible to *Mtb* infection and develop similar symptoms as humans, such as weight loss, decreased pulmonary function and the formation of caseous granulomas ^23^. As such, guinea pigs are the gold standard animal model for TB vaccine testing and are used extensively to assess the efficacy of novel TB vaccines during pre-clinical development ^24–26^. Promisingly, Δ*lsr2* vaccinated guinea pigs displayed superior protection over BCG with reduced bacterial burden, reduced organ weight and reduced lung pathology.

Δ*lsr2* is a deletion mutant strain of *Mtb* H37Rv, representing a lineage 4 strain. MTBVAC, the first live *Mtb* vaccine to begin clinical trials, is also constructed from a lineage 4 strain ^42^. There are currently nine recognized lineages (L1-9) of *Mtb*, with L2, L3, and L4 considered as modern lineages and are responsible for the majority of TB cases globally ^43,44^. In particular, L2 and L4 display increased transmissibility compared to L3 in phylogenetic studies ^44^. During development of MTBVAC, Perez et al. generated parallel deletion mutant strains from L2 and L3 to determine which construction conferred optimal vaccine potential in terms of safety and global protective efficacy. They found that the background strain lineage did not have a large impact on vaccine efficacy, as there was little to no lineage-dependent protective efficacy and good cross-lineage protection ^42^. However, the background strain lineage did have an impact on safety, with the L4 construction having the best safety profile in severe combined immunodeficient mice compared to the L2 and L3 constructions, and BCG ^42^. Thus, L4 strains appear to be the optimal background for development of live attenuated *Mtb* vaccines. This is further supported by the success of MTBVAC in Phase 1 clinical trials and its confirmed safety in adults ^12^.

The second Geneva consensus determined that live mycobacterial vaccines are a viable strategy to help control the TB epidemic, but that new *Mtb* vaccine candidates must have two stable, unmarked, independent mutations as well as show safety and efficacy at least comparable to BCG in appropriate animal models ^45^. Currently, Δ*lsr2* is PDIM deficient most likely due to spontaneous mutation ^10^. Future work will need to construct a stable PDIM mutation via an unmarked deletion of *fad26* and remove the hygromycin antibiotic-resistance marker to make Δ*lsr2* into a viable candidate to progress from the discovery stage into pre-clinical development ^17^. Further safety assessments of the stable double mutant Δ*lsr2* vaccine candidate compared to BCG in severe combined immunodeficient (SCID) mice should also be performed. SCID mice, which lack T and B lymphocytes and are highly immunocompromised, are the reference model for safety tests of live vaccines ^45,46^.

Rodent models for *Mtb* infection contain inherent limitations. Firstly, the translational capabilities of novel vaccine strategies from rodents to primates, then humans are inconsistent. This phenomenon is exaggerated in TB. This pattern is seen with *Mtb* due to the species inherent complexity and long-term co-evolution in humans ^6^. Secondly, we focused on female mice and guinea pigs for this study. Sex bias in TB incidence rates and severity is well characterized in humans, and recent work have shown these same sex biases in mice ^47,48^. As such, the observed Δ*lsr2* protection may differ in males.

A more effective vaccine to BCG would be the single greatest advantage to end the global TB epidemic ^49^. Δ*lsr2* induces a potent CD4 and CD8 T cell response and provides improved protection over BCG in both the mouse and guinea pig model of *Mtb* infection. We propose that Δ*lsr2* is a promising prototype for a future TB vaccine.

## Acknowledgements

Thank you to the excellent staff at the Division of Comparative Medicine and the Toronto High Containment Facility. We also wish to acknowledge Dr. Blair Gordon for generating the *Mtb* Δ*lsr2* mutant, Dr. Wenxi Xu for his help with flow antibody panel design, Dr. David Brooks for use of his laboratory’s flow cytometer, and Ming Li for his help with the animal work. The study was supported by Canadian Institutes for Health Research Funds PJT-156261 and PJT-186285 (J.L.). Animal figures were created with BioRender.com.

## Author contributions

Conceptualization J.W. and J.L.; Methodology J.W. and J.L.; Investigation J.W; Visualization J.W.; Funding acquisition J.L.; Project administration J.W.; Supervision J.L.; Writing – original draft J.W.; Writing – review and editing J.W. and J.L.

## Methods and Materials

### Animals

Female C57BL/6 mice between 6-10 weeks of age and female outbred Hartley guinea pigs (200-250g) were purchased form Charles River Laboratories. The high containment facility, biosafety level III laboratory at the University of Toronto, was used for all experiments involving Mycobacterium tuberculosis. The University of Toronto Animal Care Committee approved all studies and procedures were performed in accordance with the committees’ ethical standards.

### Bacterial strains and culture conditions

*Mycobacterium tuberculosis* H37Rv, *Mtb* Δ*lsr2*, and *M. bovis* BCG-Japan were prepared and titrated as previously described ^29^. Briefly, *Mtb* strains were grown at 37°C in Middlebrook 7H9 broth (BD) supplemented with 0.2% glycerol, 10% albumin-dextrose-catalase, and 0.05% Tween 80 or on 7H10 agar supplemented with 1% casamino acids (Life), 0.5% glycerol, and 10% albumin-dextrose-catalase. Hygromycin (BioShop) was added at a concentration of 75ug/mL for *Mtb* Δ*lsr2*. BCG was grown at 37°C in Middlebrook 7H9 broth (BD) supplemented with 0.2% glycerol, 10% albumin-dextrose-catalase, and 0.05% Tween 80 or on 7H10 agar supplemented with 1% casamino acids (Life), 0.5% glycerol, and 10% oleic acid-albumin-dextrose-catalase.

### C57BL/6 mice infection with mycobacterium

For aerosol inoculations, ∼100 colony-forming units (CFUs) of *Mtb* H37Rv, *Mtb* Δ*lsr2*, or *M. bovis* BCG-Japan, per mouse was delivered using a Glas-Col nebulizer Inhalation Exposure System ^29^. Inoculation dosage was confirmed by quantifying CFUs in the lungs 1 day after exposure in control mice. At various time points after infection (2, 3, 4, 5, 6, or 8 weeks) lungs and spleen CFUs were determined from homogenized tissue coated on 7H10 agar plates. CFUs were counted on the plates after a ∼4-week incubation at 37°C. A portion of the lungs and spleen were also collected for flow cytometric analysis or fixed in 10% neutral buffered formalin (Sigma Aldrich) for histological analysis.

### Intranasal vaccination of C57BL/6 mice

5×10^5^ CFU *Mtb* Δ*lsr2*, or 5×10^5^ CFU *M. bovis* BCG-Japan were delivered intranasally in 20μL PBS. 8 weeks after vaccination, mice were aerosol infected with *Mtb* as described above.

### Hartley guinea pig protection against *Mycobacterium tuberculosis* infection

Groups of 6 guinea pigs were intranasally vaccinated with 5×10^5^ CFU *Mtb* Δ*lsr2*, or 5×10^5^ CFU *M. bovis* BCG-Japan in 50uL PBS or 50uL of PBS alone as a control. At 8 weeks post-vaccination, ∼5000 CFUs of *Mtb* H37Rv per guinea pig was delivered using a Glas-Col nebulizer Inhalation Exposure System ^29^. The same batch and dilution of *Mtb* H37Rv cultures were used to infect the experimental groups under the predetermined parameter settings of the Glas-Col nebulizer. Delivery dosage was confirmed by quantifying CFU in the lungs 1 day after exposure in control guinea pigs. At 3- and 7-weeks post *Mtb* challenge, lungs and spleen CFUs were determined. Lungs and spleen were homogenized in PBS (Sigma Aldrich) and tissue homogenate was serial diluted and coated on 7H10 agar plates. CFUs were counted on the plates after a ∼4-week incubation at 37°C. A portion of the left caudal lung and spleen was fixed in 10% neutral buffered formalin (Sigma Aldrich) for histological analysis.

### Tissue isolation

Lung and spleen tissue was isolated as previously described ^50^. Briefly, lungs and spleen were chopped into fine pieces then digested in RPMI 1640 medium without calcium/magnesium (Gibco) containing 10% FBS, 1% HEPES (Life), 1 mg/mL Collagenase from Clostridium histolyticum (Sigma) and 0.15 mg/mL DNase I (Sigma) on a shaker at 37°C for 1 hour with the lungs or 30 min with the spleen. Digested tissue was filtered through a 70 μm cell strainer to obtain single-cell suspensions and an aliquot was plated on 7H10 agar plates. CFUs were counted after a ∼4-week incubation at 37°C. The remaining lung and spleen single cell suspensions were red cell-lysed and then processed for flow cytometry.

### Flow cytometry and intracellular cytokine re-stimulation

Cells were stimulated with *Mtb* lysate for 24hr and protein transport inhibitor cocktail containing Brefeldin A and Monensin was added for the final 19 hours of the stimulation time. Fixing and permeabilizing cells was done using the eBioscience Intracellular cytokine buffer kit and the eBioscience Foxp3 staining kit. All antibody dilutions and cell staining were done using PBS (Sigma-Aldrich) containing 1% fetal bovine serum (Gibco) and 2.5 mM EDTA. Zombie Aqua™ Fixable Viability Stain (BioLegend) was used to exclude dead cells from analyses.

Samples were either analyzed on FACSLyric^TM^ (BD Bioscience) in the laboratory of Dr. David Brooks or FACSSymphony^TM^ A5 (BD Bioscience) at the University of Toronto, Temerty Faculty of Medicine Flow Cytometry Core facility. Data was analyzed using Flow Jo Software v10.10 (BD FLowJo).

| **Antibodies** | **Source** | **Identifier** |
| --- | --- | --- |
| Zombie Aqua Dye | BioLegend | Cat#423101 |
| Alexa Fluor 700 anti-mouse CD45 (clone 30-F11) | BioLegend | Cat#103128; RRID: AB_493715 |
| Brilliant Violet 605 anti-mouse CD4 (clone RM4-4) | BioLegend | Cat#116027; RRID: AB_2800581 |
| Brilliant Violet 711 anti-mouse CD8a (clone 53-6.7) | BioLegend | Cat#100759; RRID: AB_2563510 |
| APC anti-mouse IFNγ (clone XMG1.2) | BioLegend | Cat#505810; RRID: AB_315404 |
| Brilliant Violet 421 anti-mouse TNFα (clone MP6-XT22) | BioLegend | Cat#506328; RRID: AB_2562902 |
| PE/Cyanine7 anti-mouse IL2 (clone JES6-5H4) | BioLegend | Cat#503832; RRID: AB_2561750 |
| PE anti-mouse IL10 (clone JES5-16E3) | BioLegend | Cat#505008; RRID: AB_315362 |
| FITC anti-mouse IL17A (clone TC11-18H10.1) | BioLegend | Cat#506907; RRID: AB_536009 |
| APC anti-mouse CD4 (clone RM4-5) | BioLegend | Cat#100515; RRID: AB_312718 |
| PE anti-mouse CD4 (clone RM4-5) | BioLegend | Cat#100511; RRID: AB_312714 |
| PE/Cyanine7 anti-mouse CD8a (clone 53-6.7) | BioLegend | Cat#100721; RRID: AB_312760 |
| Brilliant Violet 421 anti-mouse CD4 (clone GK1.5) | BioLegend | Cat#100437; RRID: AB_10900241 |
| Brilliant Violet 605 anti-mouse CD4 (clone RM4-5) | BioLegend | Cat#100548; RRID: AB_2563054 |
| Brilliant Violet 711 anti-mouse CD8a (clone 53-6.7) | BioLegend | Cat#100759; RRID: AB_2563510 |
| FITC anti-mouse CD4 (clone RM4-5) | BioLegend | Cat#100509; RRID: AB_312712 |

### Histological analysis

A portion of the left lung in mice, left caudal lung in guinea pigs and spleens from both animals were fixed for a minimum of one month in 10% neutral buffered formalin (Sigma Aldrich) and then paraffin embedded. The paraffin blocks were sectioned into 4-μm-thick slices and were stained with hematoxylin and eosin (H&E) for histological analysis. The H&E staining was performed at the Centre for Phenogenomics in Toronto.

### Quantification and statistical analysis

Graphs and statistics were generated with GraphPad Prism v10 (GraphPad Software). Number of animals per group are described in the figure legends. Error bars represent the standard error of the mean (SEM). Data shown in figures with log-scale y-axis was transformed (log_10_) to conform to normality. Student’s t-test, Mann-Whitney test, REML mixed effects model, one-way and two-way ANOVA with the Holm-Šídák or Tukey multiple comparison test were used to determine statistical significance where appropriate. Significance is indicated by: * = p < 0.05, ** = p < 0.01, *** = p < 0.001, **** = p < 0.0001, ns = not significant.

