## Supplemental Figure for "*Mycobacterium tuberculosis* Δ*lsr2* Vaccine Enhances Protection Against Tuberculosis"

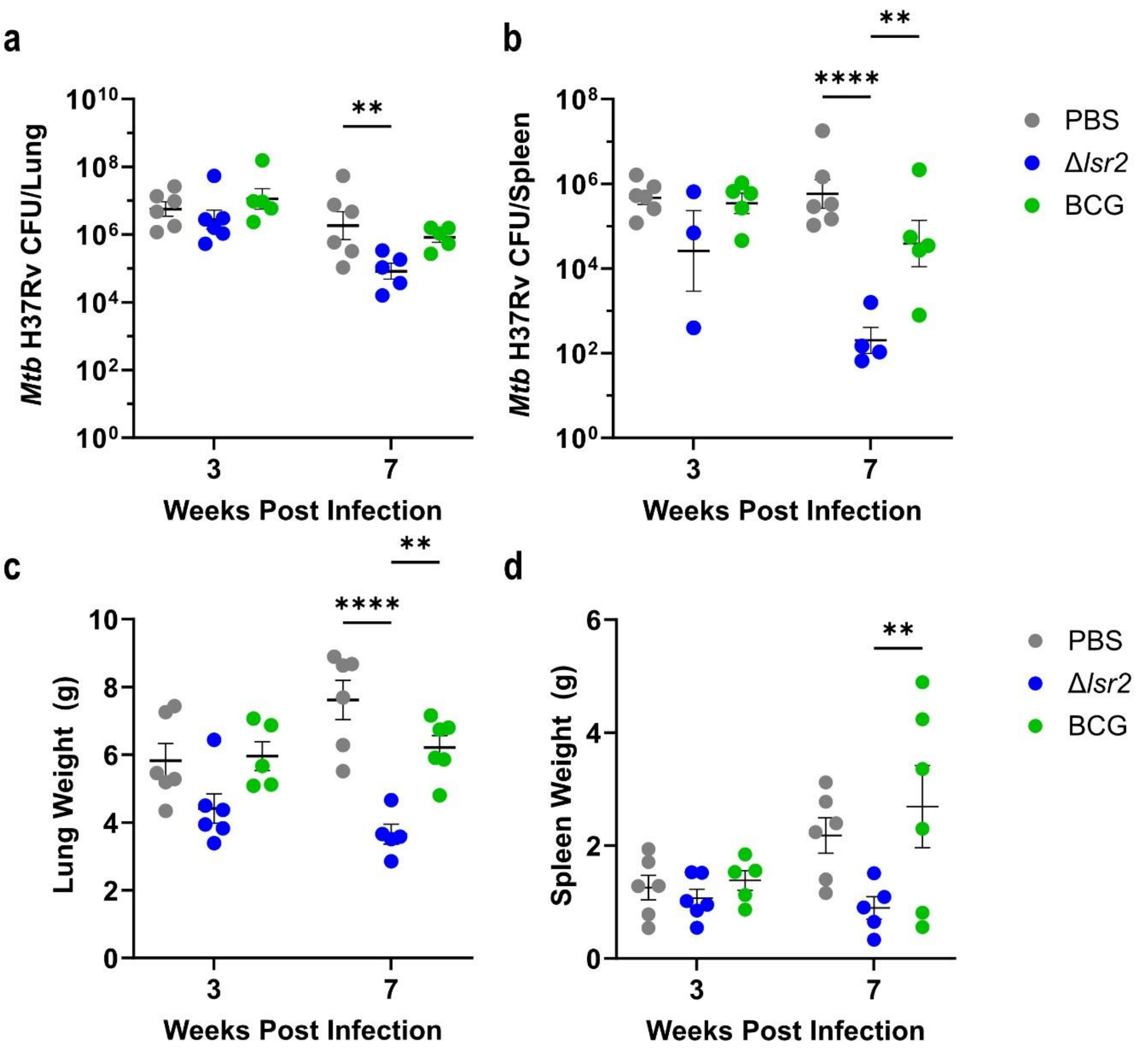


**Supplementary Figure S1 Guinea pig *Mtb* bacterial burden and organ weight without Δ*lsr2* non-responder.** a) *Mtb* H37Rv CFUs in the b) lungs and c) was determined at 3- and 7-weeks p.i. d) Lung and e) spleen organ weights were measured at 3- and 7-weeks p.i. Δ*lsr2* vaccine non-responder outlier at 7-weeks p.i. was removed. Data presented as mean ±SEM. Statistical significance was determined by two-way ANOVA with Tukey's multiple comparisons test (*, p<0.05; **, p<0.01; ***, p<0.001; ****, p<0.0001; ns = not significant).
